# No evidence for phylogenetic structure or environmental filtering of springtail microbiomes

**DOI:** 10.1101/2023.09.13.557512

**Authors:** Róbert Veres, Juliane Romahn, Clément Schneider, Miklós Bálint

## Abstract

Microorganisms play crucial roles in the lives of metazoans and can significantly impact host fitness. However, recent evidence suggests that many species may lack microorganisms that are positively associated with host fitness. Assessing the prevalence of host-specific microbiomes in animals has proven challenging due to limited studies in most higher taxa, with most investigations focusing on microbes in mammals, cephalopods, fish, and corals. This knowledge gap extends to springtails (Arthropoda: Collembola), which are widespread and abundant hexapods found in terrestrial and semi-aquatic habitats, contributing to important ecological functions. Here we investigated taxonomic bycatch in genome sequences generated from entire individuals of 70 springtail species. We aimed to understand whether microbial and other taxa associated with springtails are influenced by host phylogeny and environmental parameters. The analyses revealed high richness of bacteria and other taxa in the analyzed sequences, but detected no phylosymbiotic or environmental filtering signal in community composition. The findings suggest that springtails may be one of potentially many animal groups lacking distinct microbiomes. The study demonstrates how entire eukaryotic groups can be tested for phylosymbiotic patterns with taxonomic bycatch from genome sequences.

## Introduction

Microorganisms accompany all metazoans, and they can positively influence host fitness (Shapira 2016). Microorganisms in animals may improve nutrient uptake (Sison-Mangus, Mushegian and Ebert 2015; Visconti *et al*. 2019), contribute to resistance against pathogens (Rothacher, Ferrer-Suay and Vorburger 2016) and resilience to stress (Houtz, Taff and Vitousek 2022). Microorganisms may also influence reproductive success (Rowe *et al*. 2020) and social behaviour (Sarkar *et al*. 2020). The association with highly plastic microbiomes might even be an evolutionary strategy that allows rapid evolutionary adaptation in animals (Alberdi *et al*. 2016; Rudman *et al*. 2019). Consequently, it is increasingly common to consider hosts and associated microorganisms as holobionts (Koskella and Bergelson 2020). However, evidence is increasing that many animals lack microorganisms which are positively associated with the fitness of their hosts. A few prominent examples from diverse animal groups without a microbiome are diverse caterpillars (Hammer *et al*. 2017; Phalnikar, Kunte and Agashe 2019), ants (Hu *et al*. 2017), solitary bees (Kwong *et al*. 2017). Such animals certainly contain microbes on and in their bodies, but these are only transiently, not relevant, or sometimes detrimental to host fitness (Hammer, Sanders and Fierer 2019). As most higher animal taxa have not been studied for host-microbe associations to date (Mallott and Amato 2021), it is difficult to evaluate how wide-spread host-specific microbiomes are.

A frequent first step of screening a taxon for a potentially associated microbiome is to perform a phylosymbiotic analysis. Phylosymbiosis is defined as ‘microbial community relationships that recapitulate the phylogeny of their host’ (Brucker 2013). Such relationships were found in a many evolutionarily diverse animals. A flagship phylosymbiosis study on *Nasonia* wasps showed that gut microbiome may even contribute to host phylogenetic diversification through hybrid lethality (Brucker 2013). In vertebrates, host phylogeny and diet explains the diversity of the gut microbiome, with a particularly strong relationship between mammalian and associated microbiome phylogenies (Youngblut *et al*. 2019). A systematic study of coral reef-associated invertebrates found phylogenetic signal in all investigated taxa (O’Brien *et al*. 2020). However, the existence of phylosymbiotic pattern does not necessarily indicate tight host-microbe association (Lim and Bordenstein 2020; Mallott and Amato 2021). Such patterns may result from shared and phylogenetically conserved host traits such as food choice (Gong *et al*. 2018), and they may emerge also from methodological issues such as contamination, or difficulties to distinguish between resident and transient microbes (Hammer, Sanders and Fierer 2019). Consequently, the discovery of phylosymbiotic signal is considered as a first step for analyzing possible ecological or evolutionary processes and consequences for host-microbe systems (Lim and Bordenstein 2020).

Most phylosymbiotic analyses were performed in a few popular taxa, such as mammals, cephalopods, fish and corals (Mallott and Amato 2021). This leaves most of higher-level animal diversity not, or poorly investigated for host-microbe associations. For example, springtails (Arthropoda: Collembola) are wide-spread hexapods in terrestrial and semi-aquatic habitats, with over 9000 described species. Reaching abundances of up to 100,000 individuals per m^2^ (Coleman 2013), they contribute to important ecological functions in soils (FAO *et al*. 2020), but as for most animal groups, little information exists about their potential microbiomes (Leo *et al*. 2021). However, the presence of several interesting traits in springtails argues for further and deeper investigation of phylosymbiotic patterns. Springtails are the only known metazoans group which are able to actively synthesize antibiotics (Suring *et al*. 2017), and such ability to actively filter microbiomes is recognised as a key driver in microbiome assembly (Mazel *et al*. 2018). Springtails are also important components of underground food webs as microbivores and detritivores (Potapov *et al*. 2022), with an ability to digest complex macromolecules like cellulose and chitin (Berg, Stoffer and Heuvel 2004). The presence of endogenous cellulases (i.e. with genes encoded into the genome of an organism) was reported in several species (Busch, Danchin and Pauchet 2019). This raises the question whether and to what extent springtails may rely on their own cellulolytic abilities for obtaining nutrients from lignocellulose, in contrast to a microbiome. Here we investigated a large collection of high-throughput sequencing raw reads obtained by genome sequencing of 70 springtail species for taxonomic bycatch: sequences originating microbes and other taxa, which are not part of the targeted species’ genomes. We compared the resulting dataset to springtail phylogeny and environmental parameters, to identify potential phylosymbiotic or environmental filtering signals in microbiome composition.

## Material and methods

### Sample collection DNA Extraction and sequencing

Springtail specimens investigated here were collected to generate a taxonomically broad genome database of Central European springtails, as described in detail by (Collins *et al*. 2023). Briefly, springtail specimens of 70 species were acquired through field collection and supplemented with soil invertebrate specimens sourced from cultures and existing collections at the Senckenberg Museum (Supplementary Table 1). Sampling activities were conducted between 2011 and 2020, predominantly in Germany, although some specimens were also obtained from other European countries. Specimens were extracted from soil samples using MacFadyen or Berlese extraction. Whenever possible, non-destructive DNA extraction was performed (Gilbert *et al*. 2007). Voucher specimens have been deposited in the Görlitz collection of the Senckenberg Museum to serve as reference materials. Sequencing libraries for specimens were prepared using either the BEST protocol (Carøe *et al*. 2018) or the NEBnext ULTRA II DNA Library Prep Kit (New England Biolabs, Frankfurt am Main, Germany). The sequencing process involved short-read Illumina sequencing (300-bp paired-end), and was conducted at Novogene Europe (Cambridge, UK) on NovaSeq 6000 platforms.

**Table 1.**
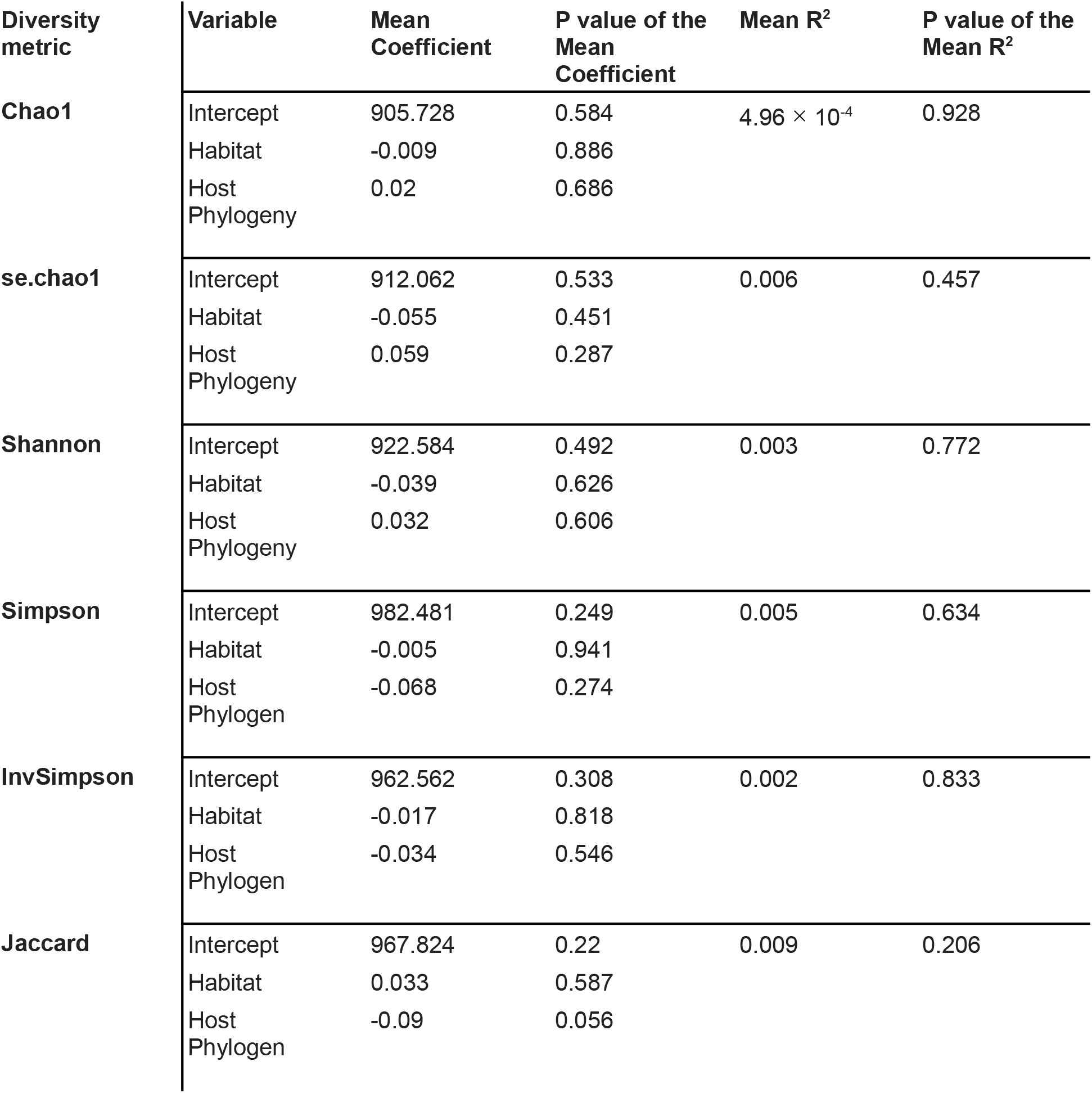
MRM model statistics for the 6 diversity metrics.

### Data Pre-processing

Adapters were trimmed with Trimmomatic (Bolger, Lohse and Usadel 2014). Sequence classification was performed using Kraken2 (Wood, Lu and Langmead 2019), a reference-based approach. Prior to classification a custom Kraken2 database was constructed, which included all available reference genomes of human, bacteria (16,572), fungi (477), plants (159), viruses (47), and protozoa (89) found on NCBI RefSeq as of January 15, 2023. In addition, the database contained 232 soil invertebrate genomes from the MetaInvert project (Collins *et al*. 2023). The inclusion of human and soil invertebrate genomes facilitated the removal of host sequences and potential human contamination from the dataset. The Kraken2 assignment results consisted of sequence counts with best possible taxonomic assignments (Suppl. Table 2), retaining bacterial, protist and viral taxa as potentially springtail-associated microbes, and plants and fungi as possible food sources.

### Statistical analyses

We estimated springtail-associated taxonomic α-(Chao1, se.chao1, Shannon, Simpson, InvSimpson) and β-diversity (Jaccard). We explained microbiome diversity with host phylogeny and habitat preferences. We derived habitat preferences of the analyzed springtails from known occurrences in different habitat types as described by level 2 hierarchies of the Coordination of Information on the Environment (CORINE, (Steemans 2008)), for more details see (Collins *et al*. 2023). As a single species may occur in multiple habitats, we constructed a table with binary columns representing each potential habitat, with values of 1 or 0 denoting the presence or absence of a particular host species in a given habitat. Subsequently, the table was transformed into a distance matrix using the ‘vegdist’ function from the vegan package (Oksanen *et al*. 2022), with Jaccard as distance metric.

Host phylogenetic tree was constructed using a multi-step approach. Firstly, two representative genomes were downloaded from the NCBI and GenBank: *Machilis hrabei* (GCA_003456935.1), the only representative of Archnaeognatha, and *Drosophila albomicans* (GCA_009650485.1), a chromosome level representative with the best contigs N50. Next, 300 genes were selected for the analysis. These genes were compared with the BUSCO arthropoda_odb10 database against fly (default) (v4.1.4) with --mode genome to ensure that the genes were conserved across the arthropod group. Genes that were present in less than 75% of the species were removed from the analysis. The remaining genes were aligned using MAFFT v7.407 (Katoh and Standley 2013) with the command ‘mafft --maxiterate 1000 –localpair’, and non-parsimony informative sites and completely conserved sites were removed using Clipkit v1.3.0 (Steenwyk *et al*. 2020). A supermatrix was then created using FASconCAT v.1.04 (Kück and Longo 2014). Finally, three gene trees were constructed using IQTree v1.6.9 (Nguyen *et al*. 2015) with 1000 bootstrap replications each. The substitution model VT+R10 was chosen according to Bayesian information criteria. The resulting trees were evaluated, and the one with the best score was selected as the final springtail phylogenetic tree.

We utilized multiple regressions on matrices (MRMs: (Lichstein 2007)) to test how well the composition of springtail-assocaited microbial communities could be explained by springtail phylogeny and host habitat type. MRM was selected as a suitable statistical method since it can be used with multiple predictors, similarly to multiple linear regression, and in contrast with simple matrix correlations (e.g. Mantel or Procrustes tests), and since both response and explanatory variables could be easily represented as distance matrices. The analyses were done with rank-based correlations. Effect size and significance were determined by comparing true data with randomly permuted data (n = 10000) in each MRM model. To assess the impact of within-species microbial variations and metadata on our MRM analysis, we conducted our analysis over 2000 randomly chosen subsample of each host species if multiple samples per species were available (Supp. Table 1). By selecting one subsample at random per host species, the variation within each species is minimized, which allows for a more comprehensive analysis of the impact of inter-species variation on overall trends and patterns. Analyzing multiple compositional scenarios in this manner provides a robust approach for excluding outlier cases that may lead to significant model results, despite underlying trends suggesting a non-significant correlation between variables. This is particularly important in cases where a minority of subsamples exhibit statistically significant model fits, while the majority do not.

Potential clustering patterns were visually examined with heatmaps and principal coordinates analysis (PCoA). PCoA analysis was performed using the ape (Paradis and Schliep 2019) and vegan (Oksanen *et al*. 2022) R packages. The heatmaps were made with the ggtree (Yu 2022), gplots (Warnes *et al*. 2022) and Heatplus (Ploner 2023) R packages, using Jaccard distances, with the color intensity in the heatmap representing the degree of dissimilarity between the microbiome compositions. The hierarchical clustering of the heatmap was performed with complete linkage, which groups microbiomes based on the maximum distance between individual objects in each cluster.

General data manipulation and basic analyses of the dataset were performed in R (v4.1.3) (R Core Team 2023) with the vegan, phyloseq (McMurdie and Holmes 2013), dplyr, tidyr, and ggplot2 R packages. Alpha and Beta diversity were calculated with the phyloseq R package. MRMs were performed with the ecodist (Goslee and Urban 2007) R package, using scripts adapted from (Youngblut *et al*. 2019).

## Results

### Taxonomic richness and composition

A total of approximately 2.83 × 10^9^ sequences from 78 springtail samples was available for the analysis. The Kraken2 search of these sequences revealed that 74.27% of the sequences were classified as soil invertebrates, 1% as bacteria, 0.82% as human, 0.01% as fungi, 0.01% as plant, < 0.001% as viral, < 0.001% as protozoa, and 23.89% could not be identified (Supp. Table 2). The human, soil invertebrate and unidentified sequences were removed from the data set, resulting in approximately 2.86 × 10^7^ sequences. The final data was composed of bacterial (98.81%), fungal (0.59%), plant (0.58%), viral (<0.001%), and protozoan (0.02%) sequences (Fig 1a), belonging to 6409 bacterial, 322 fungal, 223 plant, 6 viral and 40 protozoan taxa. A total of 27 bacterial phyla were identified, of which Proteobacteria were the most taxonomically rich (accounting for 45% of all identified bacterial taxa), followed by Actinomycetota (21%) and Bacillota (17%) (Fig 1b).

**Fig. 1.**
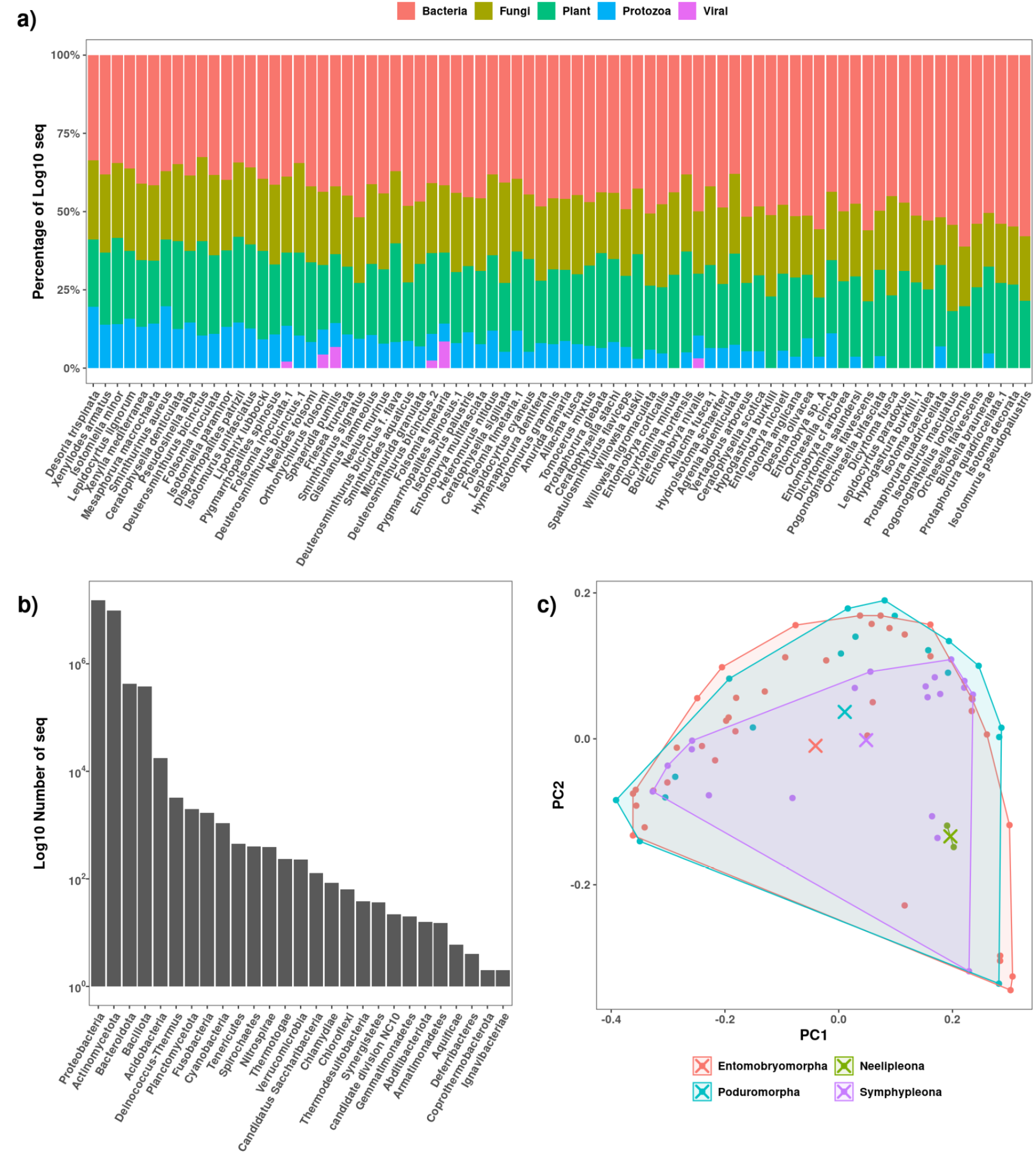
**a)**. Per-domain sequence ratios of non-springtail reads in 70 springtail hosts. Abundances were log-transformed before calculating the percentage composition. **b)**. Log-transformed read abundances of the identified bacterial phyla. **c)**. Principal Coordinate Analysis of the host microbiomes at the order level, with PC1 and PC2 explaining 16.80% and 5.89% of variation in Jaccard distances.

### Predictors of community composition

When analyzing all bycatch together, none of the six MRM models, each explaining a different diversity metric of springtail-associated taxa with host phylogeny and habitat preferences, showed a statistically significant overall fit. This finding was further corroborated by the principal coordinate analysis (PCoA) ordinations of beta diversity values (Fig. 1c), and a heatmap visualization of taxon distributions across the host phylogeny (Fig. 2), with none of these showing discernible clustering by host phylogeny. When analysing bacteria, fungi and plants separately, MRM models of bacterial diversity metrics did not show any statistically significant fit (Supp. Table 3). MRM models of fungi and plants showed a weak, but statistically significant decrease in community distance with increasing host phylogenetic distance (Supp. Tables 4-5). The visual inspection of heatmaps hinted at randomly assembled bacterial, fungal and plant communities (Supp. Fig. 1).

**Fig. 2.**
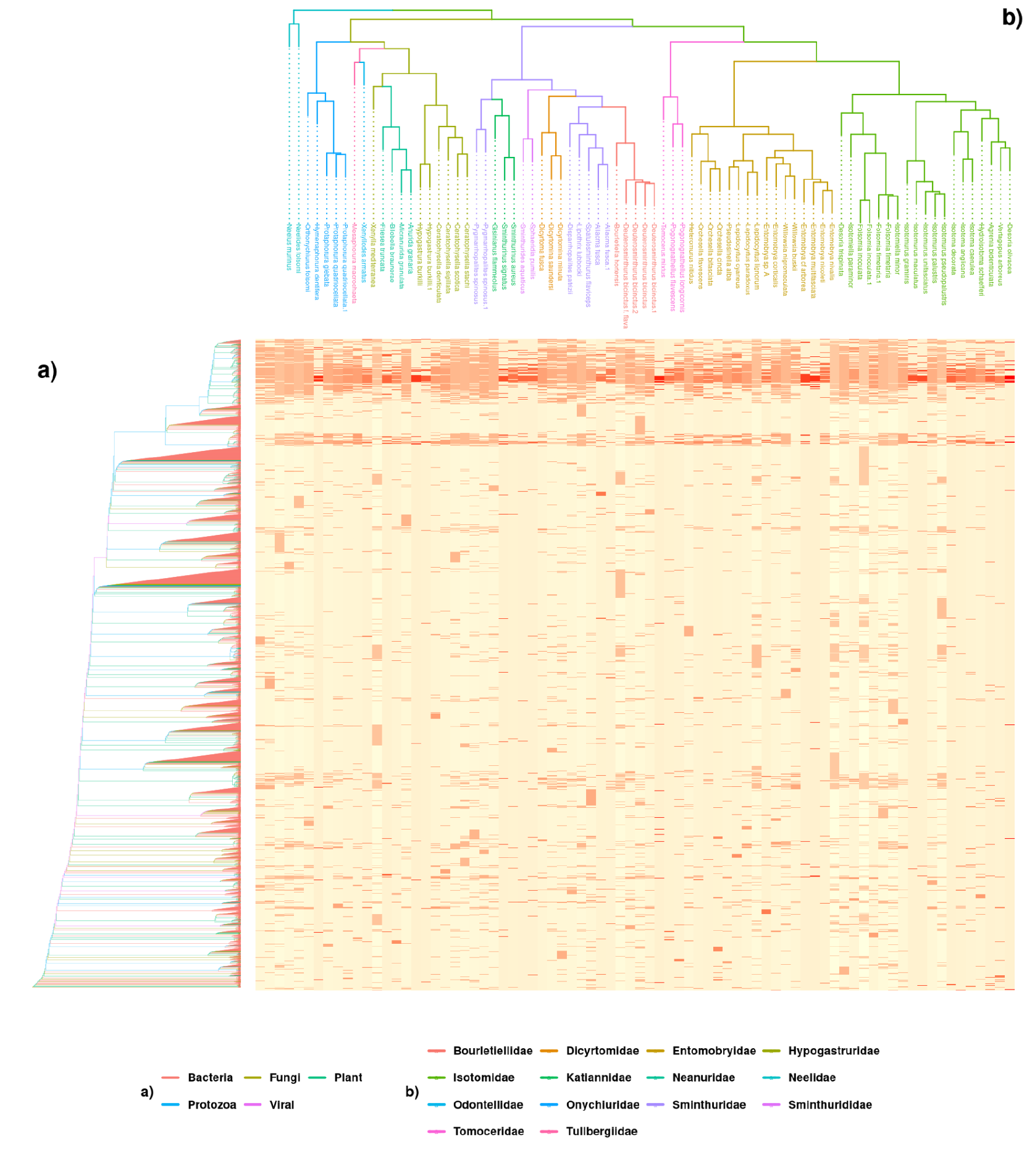
Relationship between taxonomic bycatch from springtail genome sequences and the phylogeny of springtails. The X-axis represents phylogenetic relationships among 70 springtail species. (b). The Y-axis represents microbiome OTUs categorized into bacteria, fungi, plants, viruses, and protozoa (a). The heatmap is color-coded to represent the relative abundance of each microbiome OTU in each of the soil invertebrate species. The intensity of the color corresponds to the degree of association, with darker shades indicating higher abundance.

## Discussion

### Taxonomic diversity of springtail microbiomes

We identified sequences from diverse bacterial (6409 taxa) and fungal communities (322 taxa) in springtails, along with plants (223 taxa), and some viruses and protists. The most taxon-rich higher groups of bacteria were Proteobacteria, Actinomycetota and Bacillota, reflecting what was found in Antarctic springtail microbiomes (Leo *et al*. 2021). This is on the same order of magnitude as reported before on marker-gene-based OTUs of bacteria (Leo *et al*. 2021). However, fungal richness reported here is about one order of magnitude lower than previous reports from springtails based on amplified marker gene OTUs (Anslan, Bahram and Tedersoo 2018). These differences might stem from methodological differences: while OTUs in metabarcoding studies such as (Anslan, Bahram and Tedersoo 2018; Leo *et al*. 2021) rely on OTUs generated by clustering PCR-amplified marker gene sequences, we identified taxonomic bycatch from genome sequences. These differences might be compounded by differences in the availability of sequence databases which can be used for taxonomic identification. Our sequence identification is based on 16,572 bacterial, but only on 477 fungal publicly available genomes. This contrasts with the availability of marker gene reference sequences: the SILVA database (release 138) contains around 123,000 bacterial 16S sequences (Quast *et al*. 2013), while the UNITE database (v.9.0) contains over 8,390,000 ITS sequences, corresponding to over 290,000 fungal species hypothesis (Nilsson *et al*. 2019).

### No effect of host phylogeny or habitat preference on the microbiome

We did not find a statistically significant association between bacterial communities of springtails, host phylogeny, and the environment. Host phylogeny and environmental factors together explained only ∼1% of variation in different aspects of microbiome community composition and diversity (Supp. Table 3). The lack of statistically significant phylosymbiotic signal confirms previous findings that microbial communities of springtails only weakly reflect taxonomic differences among hosts (Leo *et al*. 2021). Interestingly, we could not detect a recently reported phylogenetic signal in Wolbachia infections in springtails (Rodrigues *et al*. 2023), despite the identification of several Wolbachia strains in the sequence data (Supp. Table 2). The lack of environmental signal contrasts with what was found in the microbial communities of four Antarctic springtails (Leo *et al*. 2021). However, as Leo et al. point out, environmental filtering of microbes in that study has to be carefully interpreted as one species was reared under laboratory conditions for 18 months before analysis.

Even though animal-associated microbiomes were found in numerous and taxonomically diverse study systems, animals that lack associated microbiomes also seem to be common (Hammer, Sanders and Fierer 2019). Although we are far from having comparative studies for most animal classes (Mallott and Amato 2021), there are prominent examples where animals do not rely on fitness benefits provided by microbiomes. For example, bacteria seem to be transitory in caterpillar guts where they occur in low abundances, likely originate from plant food, and do not seem to influence several fitness parameters (Phalnikar, Kunte and Agashe 2019). Microbiomes of microscoping marine invertebrates from 21 phyla do not reflect host phylogeny, but rather environmental effects (Boscaro *et al*. 2022). Even in vertebrates, which often harbor specific microbiomes, gut microbes of birds and bats are only weakly related to host phylogeny or diet (Song *et al*. 2020).

Animals without microbiomes miss out on benefits that associated microbes may confer, but they also do not pay associated costs, such as competition for nutrients, symbiont potential for parasitism of pathogenicity, increased sensitivity to environmental perturbations (reviewed in (Hammer, Sanders and Fierer 2019). A well-studied function conferred by microbiomes to hosts is defense against pathogens (Rothacher, Ferrer-Suay and Vorburger 2016). However, many springtails are able to synthesize their own antibiotics (Suring *et al*. 2017), so they might not need protective microbiomes. Another well-studied function of microbiomes is contribution to host metabolism (Sison-Mangus, Mushegian and Ebert 2015; Visconti *et al*. 2019). However, some studies indicate that at least several springtails are capable of the endogenous production of cellulases (Hong *et al*. 2014; Busch, Danchin and Pauchet 2019). Springtails thus might directly access plant nutrients without support from microbes, similarly to other animals that rely entirely on endogenous cellulases for lignocellulose digestion, such as several species of crustaceans (King *et al*. 2010; Kern *et al*. 2013). Phylosymbiosis also appears to be less common in animal clades with highly diverse microbial communities (Mallott and Amato 2021), and the communities investigated here with 6409 bacterial taxa were fairly diverse. Independence from microorganisms may be a successful evolutionary strategy in animals (Hammer, Sanders and Fierer 2019), and springtails seem to be one of the potentially many animal groups that do not rely on associated microbiomes. However, it is important to highlight that inability to detect a pattern is no proof about the inexistence of this pattern. Although we were able to detect several strains of *Wolbachia* (Supp. Table 2), we could not confirm recently reported phylogenetic signals in *Wolbachia* infections in springtails (Rodrigues *et al*. 2023). We suspect that the difference stems from the sensitivity of the applied methods: while Rodrigues and colleagues screened for *Wolbachia* with specific markers, our analysis relies on metagenomic shotgun sequences. This might not be sensitive enough to pick up a *Wolbachia*-specific signal among the thousands of microbial strains also detected by the shotgun approach.

### Little taxonomic differences in potential fungal and plant food sources

We identified altogether 223 plant and 322 fungal taxa in the samples, which we interpret as potential signatures of food remains in springtail guts. Springtails feed on diverse food types, being able to digest complex sugars such as chitin and cellulose (Berg, Stoffer and Heuvel 2004), with fungal hyphae and plant materials dominating their guts (reviewed by (Potapov *et al*. 2022)). We detected a very weak decrease in plant and fungal community composition with increasing phylogenetic distance among the hosts, which contrasts with isotope analyses that show phylogenetic conservatism in food choice among orders (Potapov *et al*. 2016). However, this decrease has to be interpreted with much caution considering poor model fit (Supp. Tables 4-5) and the large variation in fungal and plant community composition (Suppl. Fig. 1). Variation in fungal community composition found here is similar to previously reported high spatial and temporal variation in springtail-associated fungi (Anslan, Bahram and Tedersoo 2018). A lack of positive phylogenetic signal in plant and fungal sequences might be related to a high trophic flexibility of springtails (Potapov, Tiunov and Scheu 2019). However, it is also possible that molecular studies record DNA sequences from many fungi and plants which actually do not contribute to springtail metabolism.

### Outlook

There are several issues with our study design that need improvement in the future. First, the analyses were done on the level of best possible taxonomic assignment of reads, and this might lump together otherwise distinct microbial taxa. Such low levels of divergence among microbial strains might be resolved by marker-gene-based confirmation studies. Second, here we analyzed the full microbiome, and this may hide phylosymbiotic or environmental differences in particular body parts. Third, since we were working with taxonomic bycatch from genome sequencing of the host species, our study has no explicit experimental design to test for environmental influences. Fourth, our coverage of the springtail taxonomy is not even, with one of the orders (Neelipleona) being present with only two species in the analysis. This group with debated taxonomic position is notoriously difficult to work with: they are small even among springtails and much less diversified than the other orders. Nonetheless, their cryptic lifestyle is connected to a distinct ecology. Fifth, our results about the lack of microbiomes should be confirmed by experimental tests of fitness benefits.

Genome sequences covering all eukaryotic families are increasingly becoming available through large genome sequencing initiatives, such as the Earth BioGenome Project (Lewin *et al*. 2022) or the European Reference Genome Atlas (Formenti *et al*. 2022). Co-sequencing host species and potentially associated microbes will help identifying phylosymbiotic patterns, providing fundamental information about the prevalence of potential host-microbe associations across the tree of life. Co-sequencing hosts and microbes will also allow us to ask mechanistic questions about coexistence. In the case of springtails, such questions should address differences among microorganism communities in taxa with antibiotics synthesis capability, in comparison to taxa that cannot synthesize their own antibiotics. A similar question should address differences among springtails with respect to the presence of endogeneous cellulases. Genome annotation for a wider taxonomic range of potential hosts needs to improve to address such questions. Our study demonstrates how the analysis of taxonomic bycatch in a taxonomically comprehensive and dense sampling of genomes provides the statistical power to evaluate phylosymbiotic patterns in entire eukaryotic groups.

## Acknowledgements

The authors thank Ricarda Lehmitz, Ulrich Burkhardt, Karin Hohberg, Peter Decker, Jörg Müller for Sampling and laboratory support, and Nick Tobias for initial discussions about the analyses.

## References

Alberdi A, Aizpurua O, Bohmann K et al. Do Vertebrate Gut Metagenomes Confer Rapid Ecological Adaptation? Trends Ecol Evol 2016;31:689–99.

Anslan S, Bahram M, Tedersoo L. Seasonal and annual variation in fungal communities associated with epigeic springtails (Collembola spp.) in boreal forests. Soil Biol Biochem 2018;116:245–52.

Berg M, Stoffer M, Heuvel H. Feeding guilds in Collembola based on digestive enzymes. Pedobiologia 2004;48:589–601.

Bolger AM, Lohse M, Usadel B. Trimmomatic: a flexible trimmer for Illumina sequence data. Bioinformatics 2014;30:2114–20.

Boscaro V, Holt CC, Van Steenkiste NWL et al. Microbiomes of microscopic marine invertebrates do not reveal signatures of phylosymbiosis. Nat Microbiol 2022;7:810–9.

Busch A, Danchin EGJ, Pauchet Y. Functional diversification of horizontally acquired glycoside hydrolase family 45 (GH45) proteins in Phytophaga beetles. BMC Evol Biol 2019;19:100.

Carøe C, Gopalakrishnan S, Vinner L et al. Single-tube library preparation for degraded DNA. Methods Ecol Evol 2018;9:410–9.

Coleman DC. Soil Biota, Soil Systems, and Processes. In: Levin SA (ed.). Encyclopedia of Biodiversity (Second Edition). Waltham: Academic Press, 2013, 580–9.

Collins G, Schneider C, Boštjančić LL et al. MetaInvert: A New Soil Invertebrate Genome Resource Provides Insights into below-Ground Biodiversity and Evolution. In Review, 2023.

FAO, ITPS, GSBI et al. State of Knowledge of Soil Biodiversity - Status, Challenges and Potentialities, Report 2020. Rome, Italy: FAO, 2020.

Formenti G, Theissinger K, Fernandes C et al. The era of reference genomes in conservation genomics. Trends Ecol Evol 2022;37:197–202.

Gilbert MTP, Moore W, Melchior L et al. DNA extraction from dry museum beetles without conferring external morphological damage. PloS One 2007;2:e272.

Gong X, Chen T-W, Zieger SL et al. Phylogenetic and trophic determinants of gut microbiota in soil oribatid mites. Soil Biol Biochem 2018;123:155–64.

Goslee SC, Urban DL. The ecodist Package for Dissimilarity-based Analysis of Ecological Data. J Stat Softw 2007;22:1–19.

Hammer TJ, Janzen DH, Hallwachs W et al. Caterpillars lack a resident gut microbiome. Proc Natl Acad Sci 2017;114:9641–6.

Hammer TJ, Sanders JG, Fierer N. Not all animals need a microbiome. FEMS Microbiol Lett 2019;366:fnz117.

Hong SM, Sung HS, Kang MH et al. Characterization of Cryptopygus antarcticus Endo-β-1,4-Glucanase from Bombyx mori Expression Systems. Mol Biotechnol 2014;56:878–89.

Houtz JL, Taff CC, Vitousek MN. Gut Microbiome as a Mediator of Stress Resilience: A Reactive Scope Model Framework. Integr Comp Biol 2022;62:41–57.

Hu Y, Holway DA, Łukasik P et al. By their own devices: invasive Argentine ants have shifted diet without clear aid from symbiotic microbes. Mol Ecol 2017;26:1608–30.

Katoh K, Standley DM. MAFFT Multiple Sequence Alignment Software Version 7: Improvements in Performance and Usability. Mol Biol Evol 2013;30:772–80.

Kern M, McGeehan JE, Streeter SD et al. Structural characterization of a unique marine animal family 7 cellobiohydrolase suggests a mechanism of cellulase salt tolerance. Proc Natl Acad Sci 2013;110:10189–94.

King AJ, Cragg SM, Li Y et al. Molecular insight into lignocellulose digestion by a marine isopod in the absence of gut microbes. Proc Natl Acad Sci 2010;107:5345–50.

Koskella B, Bergelson J. The study of host–microbiome (co)evolution across levels of selection. Philos Trans R Soc B Biol Sci 2020;375:20190604.

Kück P, Longo GC. FASconCAT-G: extensive functions for multiple sequence alignment preparations concerning phylogenetic studies. Front Zool 2014;11:81.

Kwong WK, Medina LA, Koch H et al. Dynamic microbiome evolution in social bees. Sci Adv 2017;3:e1600513.

Leo C, Nardi F, Cucini C et al. Evidence for strong environmental control on bacterial microbiomes of Antarctic springtails. Sci Rep 2021;11:2973.

Lewin HA, Richards S, Lieberman Aiden E et al. The Earth BioGenome Project 2020: Starting the clock. Proc Natl Acad Sci 2022;119:e2115635118.

Lichstein JW. Multiple regression on distance matrices: a multivariate spatial analysis tool. Plant Ecol 2007;188:117–31.

Lim SJ, Bordenstein SR. An introduction to phylosymbiosis. Proc R Soc B Biol Sci 2020;287:20192900.

Mallott EK, Amato KR. Host specificity of the gut microbiome. Nat Rev Microbiol 2021;19:639–53.

Mazel F, Davis KM, Loudon A et al. Is Host Filtering the Main Driver of Phylosymbiosis across the Tree of Life? mSystems 2018;3:e00097–18.

McMurdie PJ, Holmes S. phyloseq: An R Package for Reproducible Interactive Analysis and Graphics of Microbiome Census Data. PLOS ONE 2013;8:e61217.

Nguyen L-T, Schmidt HA, von Haeseler A et al. IQ-TREE: A Fast and Effective Stochastic Algorithm for Estimating Maximum-Likelihood Phylogenies. Mol Biol Evol 2015;32:268–74.

Nilsson RH, Larsson K-H, Taylor AFS et al. The UNITE database for molecular identification of fungi: handling dark taxa and parallel taxonomic classifications. Nucleic Acids Res 2019;47:D259–64.

O’Brien PA, Tan S, Yang C et al. Diverse coral reef invertebrates exhibit patterns of phylosymbiosis. ISME J 2020;14:2211–22.

Oksanen J, Simpson GL, Blanchet FG et al. Vegan: Community Ecology Package., 2022.

Paradis E, Schliep K. ape 5.0: an environment for modern phylogenetics and evolutionary analyses in R. Bioinformatics 2019;35:526–8.

Phalnikar K, Kunte K, Agashe D. Disrupting butterfly caterpillar microbiomes does not impact their survival and development. Proc R Soc B Biol Sci 2019;286:20192438.

Ploner A. Heatplus: Heatmaps with row and/or column covariates and colored clusters. R package version 3.8.0. 2023.

Potapov AA, Semenina EE, Korotkevich AYu et al. Connecting taxonomy and ecology: Trophic niches of collembolans as related to taxonomic identity and life forms. Soil Biol Biochem 2016;101:20–31.

Potapov AM, Beaulieu F, Birkhofer K et al. Feeding habits and multifunctional classification of soil-associated consumers from protists to vertebrates. Biol Rev 2022;97:1057–117.

Potapov AM, Tiunov AV, Scheu S. Uncovering trophic positions and food resources of soil animals using bulk natural stable isotope composition. Biol Rev 2019;94:37–59.

Quast C, Pruesse E, Yilmaz P et al. The SILVA ribosomal RNA gene database project: improved data processing and web-based tools. Nucleic Acids Res 2013;41:D590–6.

R Core Team. R: A Language and Environment for Statistical Computing. Vienna, Austria: R Foundation for Statistical Computing, 2023.

Rodrigues J, Lefoulon E, Gavotte L et al. Wolbachia springs eternal: symbiosis in Collembola is associated with host ecology. R Soc Open Sci 2023;10:230288.

Rothacher L, Ferrer-Suay M, Vorburger C. Bacterial endosymbionts protect aphids in the field and alter parasitoid community composition. Ecology 2016;97:1712–23.

Rowe M, Veerus L, Trosvik P et al. The Reproductive Microbiome: An Emerging Driver of Sexual Selection, Sexual Conflict, Mating Systems, and Reproductive Isolation. Trends Ecol Evol 2020;35:220–34.

Rudman SM, Greenblum S, Hughes RC et al. Microbiome composition shapes rapid genomic adaptation of Drosophila melanogaster. Proc Natl Acad Sci 2019;116:20025–32.

Sarkar A, Harty S, Johnson KV-A et al. The role of the microbiome in the neurobiology of social behaviour. Biol Rev 2020;95:1131–66.

Shapira M. Gut Microbiotas and Host Evolution: Scaling Up Symbiosis. Trends Ecol Evol 2016;31:539–49.

Sison-Mangus MP, Mushegian AA, Ebert D. Water fleas require microbiota for survival, growth and reproduction. ISME J 2015;9:59–67.

Song SJ, Sanders JG, Delsuc F et al. Comparative Analyses of Vertebrate Gut Microbiomes Reveal Convergence between Birds and Bats. mBio 2020;11:e02901–19.

Steemans C. Coordination of Information on the Environment (CORINE). Encycl Geogr Inf Sci Ed Kemp K Sage Publ Inc Thousand Oaks CA 2008:49–50.

Steenwyk JL, Iii TJB, Li Y et al. ClipKIT: A multiple sequence alignment trimming software for accurate phylogenomic inference. PLOS Biol 2020;18:e3001007.

Suring W, Meusemann K, Blanke A et al. Evolutionary ecology of beta-lactam gene clusters in animals. Mol Ecol 2017;26:3217–29.

Visconti A, Le Roy CI, Rosa F et al. Interplay between the human gut microbiome and host metabolism. Nat Commun 2019;10:4505.

Warnes GR, Bolker B, Bonebakker L et al. Gplots: Various R Programming Tools for Plotting Data., 2022.

Wood DE, Lu J, Langmead B. Improved metagenomic analysis with Kraken 2. Genome Biol 2019;20:257.

Youngblut ND, Reischer GH, Walters W et al. Host diet and evolutionary history explain different aspects of gut microbiome diversity among vertebrate clades. Nat Commun 2019;10:2200.

Yu G. Data Integration, Manipulation and Visualization of Phylogenetic Trees. New York: Chapman and Hall/CRC, 2022.

