## Supplementary Figure 1 for "No evidence for phylogenetic structure or environmental filtering of springtail microbiomes"

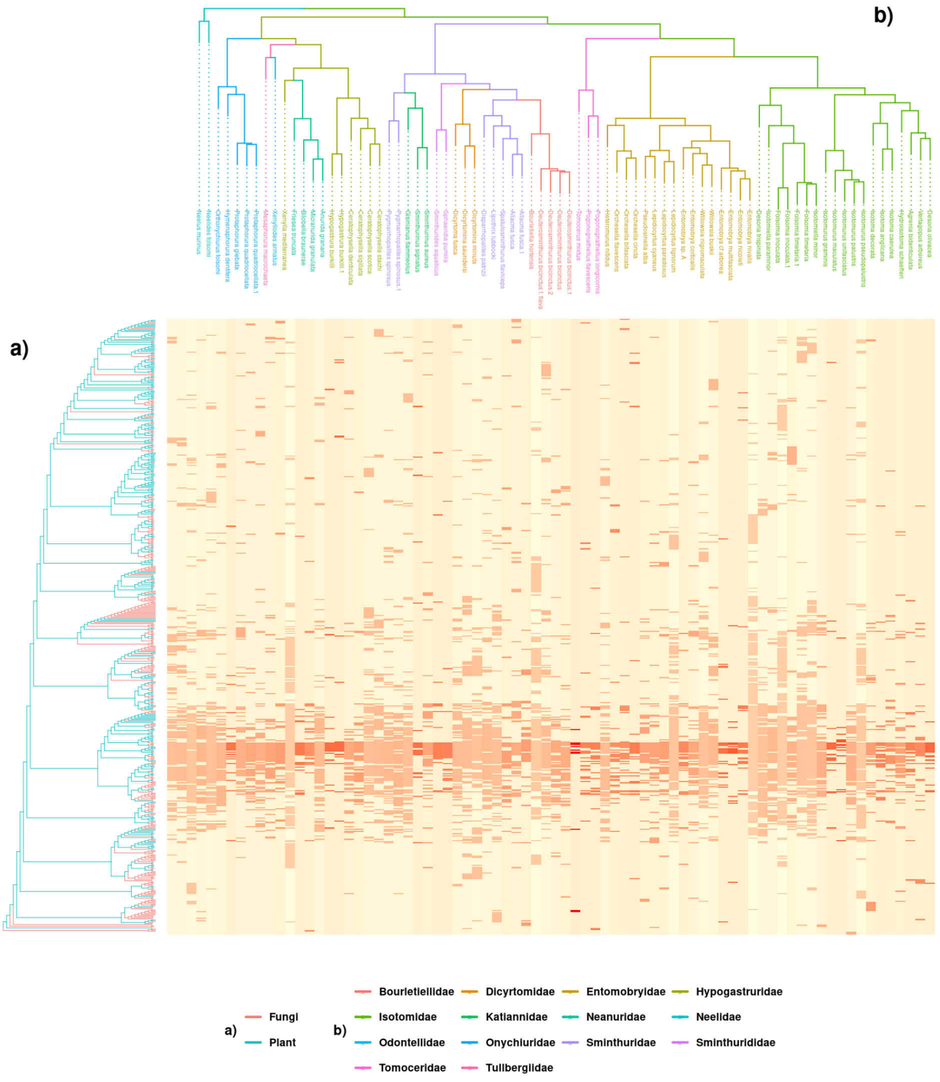


Supp. Fig. 1. Relationship between fungal and plant taxa and the phylogeny of springtails. The X-axis represents phylogenetic relationships among 70 springtail species. The heatmap is color-coded to represent the relative abundance of each microbiome OTU in each of the soil invertebrate species. The intensity of the color corresponds to the degree of association, with darker shades indicating higher abundance.
