## Supplementary Table Legends for "No evidence for phylogenetic structure or environmental filtering of springtail microbiomes"

### Supplementary Tables

Supp. Table 1. Collection and habitat details, and sequence availability of springtail host specimens

Supp. Table 2. Taxonomic assignments of bacterial, fungal, plant, protist and viral sequences co-sequenced with springtail hosts

Supp. Table 3. MRM model results of six diversity metrics of bacteria

Supp. Table 4. MRM model results of six diversity metrics of plants

Supp. Table 5. MRM model results of six diversity metrics of fungi
